# A minimal model demonstrates the control of mouse sperm capacitation by a positive feedback loop between sNHE and SLO3

**DOI:** 10.1101/2021.11.30.469438

**Authors:** Bertrand de Prelle, Pascale Lybaert, David Gall

**Affiliations:** Research Laboratory on Human Reproduction, Faculté de Médecine, Université libre de Bruxelles, Belgium

**Keywords:** sperm, capacitation, sNHE, SLO3, hyperpolarization, mathematical model, bistability

## Abstract

When mammalian spermatozoa are released in the female reproductive tract, they are incapable of fertilizing the oocyte. They need a prolonged exposure to the alkaline medium of the female genital tract before their flagellum gets hyperactivated and the acrosome reaction can take place, allowing the sperm to interact with the oocyte. Ionic fluxes across the sperm membrane are involved in two essential aspects of capacitation: the increase in intracellular pH and the membrane hyperpolarization. In particular, it has been shown that the SLO3 potassium channel and the sNHE sodium-proton transporter, two sperm-specific transmembrane proteins, are necessary for the capacitation process to occur. As the SLO3 channel is activated by an increase in intracellular pH and sNHE is activated by hyperpolarization, they act together as a positive feedback system. Mathematical modeling provides a unique tool to capture the essence of a molecular mechanism and can be used to derive insight from the existing data. We have therefore developed a theoretical model formalizing the positive feedback loop between SLO3 and sHNE in mouse epididymal sperm to see if this non-linear interaction can provide the core mechanism explaining the existence of uncapacited and capacitated states. We show that the proposed model can fully explain the switch between the uncapacitated and capacited states and also predicts the existence of a bistable behaviour. Furthermore, our model indicates that SLO3 inhibition, above a certain threshold, is effective to completely abolish capacitation.

## 1 INTRODUCTION

Despite continuous research in reproductive biology over the last two decades, the prevalence of couple infertility (over 12 months) remains around 15%, among which 30% is due to a male infertility factor (Hajder et al., 2016). The sperm count has been continuously decreasing for 40 years, raising the alarm for a major fertility crisis by the midst of the 21st century and the need for increased research in male infertility(Levine et al., 2017; Barratt et al., 2017; Duffy et al., 2020). Our knowledge of the molecular regulation of sperm motility and its fertilization potential is still incomplete and the etiology of a number of human male infertility cases remains unknown. Therefore, whether in search of new male fertility screening methods or novel contraceptive solutions, a deeper understanding of the molecular events regulating sperm functions is needed. These functions notably depend on ion homeostasis, which is controlled by ion channels and transporters. Many of these proteins or their regulatory subunits are expressed exclusively in sperm cells, making them ideal pharmacological targets (Wang et al., 2021).

Before mammalian spermatozoa are able to to fertilize the oocyte, they need to spend some time in the female genital tract. In human this duration must be of several hours while in mouse it is around an hour. During this transit, sperms are exposed to a diversity of environmental and intracellular signals allowing sperm to acquire a special form of motility, known as hyperactivation, and the ability to undergo the acrosome reaction. This process is called capacitation and since its discovery (Austin, 1951; Chang, 1951), *in vitro* studies of mammalian sperm showed that the presence of albumin and bicarbonate in the physiological incubation medium is essential for capacitation to occur (Lee and Storey, 1986; Stival et al., 2015).

Mammalian sperm capacitation is characterized by an increase in intracellular pH (pH_i_) (Zeng et al., 1996), membrane hyperpolarization (Arnoult et al., 1999) and a calcium influx from the extracellular medium (Ruknudin and Silver, 1990), and since nearly three decades now, pharmacological and genetic studies have revealed the presence of sperm-specific proteins that are essential for male fertility and necessary for the sperm to reach capacitation: the sNHE sodium-proton exchanger, the SLO3 potassium channel and the CATSPER calcium channel. The absence of any of the corresponding genes leads to male infertility without any systemic abnormality, in accordance with the fact that these proteins are expressed only in the sperm (Wang et al., 2003; Santi et al., 2010; Ren et al., 2001).

The calcium influx, increase in pH_i_ and hyperpolarization observed during capacitation are dependent on the activity of CATSPER, sNHE and SLO3 (Carlson et al., 2003; Wang et al., 2003; Santi et al., 2010), and these actors operate all together as CATSPER and SLO3 are activated by an elevation of pH_i_ and sNHE is activated by hyperpolarization (Kirichok et al., 2006; Schreiber et al., 1998; Windler et al., 2018). These considerations led Chávez et al. (2014) to the proposal of the existence of a positive feedback loop between the activation of sNHE and SLO3, leading to a high pH_i_ and membrane hyperpolarization, that could promote the pH_i_-dependent CATSPER’s activity during capacitation.

In order to test this hypothesis and find if capacitation can indeed be controlled by the feedback loop between sNHE and SLO3, we propose a mathematical model for the capacitation process based on these two essential molecular actors of capacitation. Using this minimal model we then investigate the conditions necessary for the incubation medium to capacitate sperms and ask whether this process of capacitation can be reversed back. Finally we also investigate the most effective way of preventing capacitation by inhibition of these two proteins.

## 2 MATERIALS AND METHODS

Our model is a minimal two variables model describing the evolution in time of the intracellular pH (pH_i_) and the transmembrane electrical potential (*V*_m_) of a mouse epididymal spermatozoon, which includes the feedback between the increase in pH_i_ and hyperpolarization resulting from the activations of sNHE and SLO3.

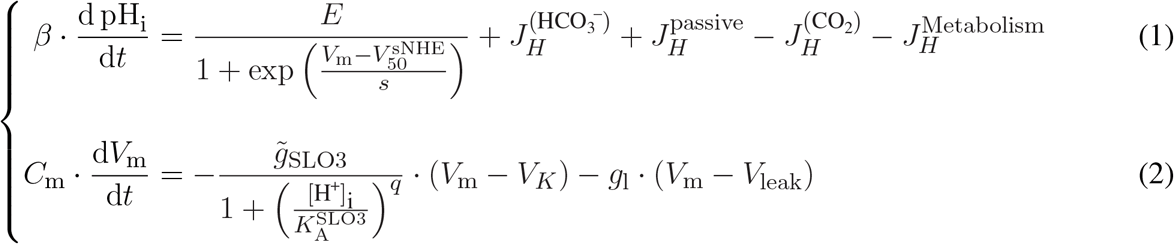

The first equation (1) links the evolution of pH_i_ to the different mechanisms related to proton homeostasis (figure 1A), where the *β* factor is the total pH_i_ buffer capacity of the sperm cell. The second equation (2) is based on the conservation of the electrical charge and shows that the evolution of the transmembrane potential depends on the contributions of SLO3 and a leak current. The factor *C*_*m*_ is the membrane’s capacitance of the sperm cell. All the actors taken into account in the two equations of the model are schematized on figure 1A.

**Figure 1.**
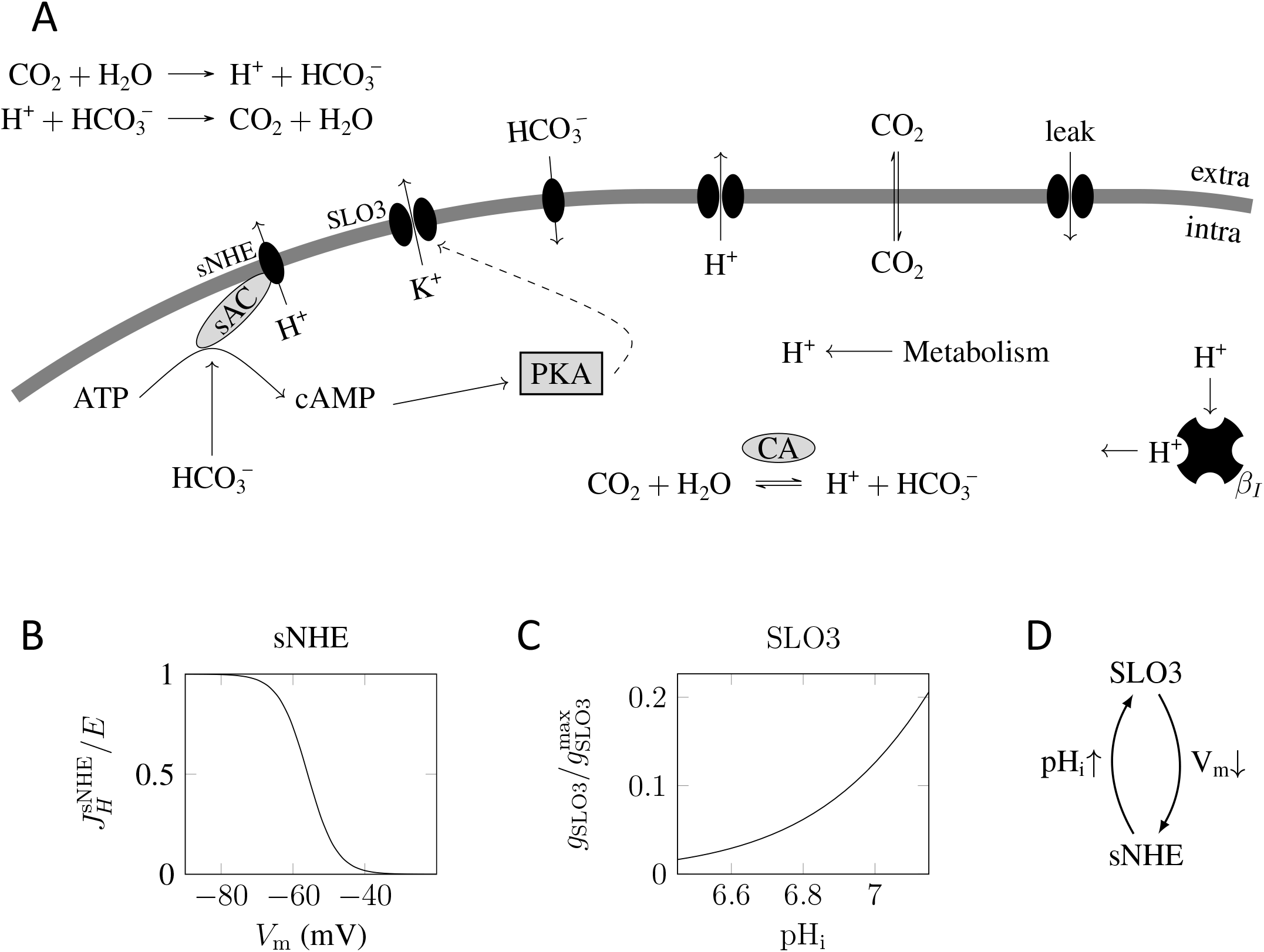
The feedback loop regulating capacitation. (A) The model takes into account the potassium channel SLO3 and the sodium-proton exchanger sNHE. SLO3 is activated downstream of cAMP production through the activation of PKA. The cAMP production by the soluble adenylyl cyclase is activated by the presence of bicarbonate 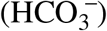. A leak current, regrouping all transmembrane electrical currents except through SLO3, is taken into account. 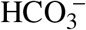 passes through the plasma membrane by a variety of transporters. The carbon dioxyde (CO_2_) freely and instantly crosses the cell membrane, and equilibrates with 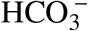 and H^+^. An influx of protons resulting from the metabolism is taken into account and the intracellular pH varies due to a leak of protons through the plasma membrane. Finally the total intrinsic protons buffer β_*I*_ is present. (B) sNHE activation curve against *V*_m_. Hyperpolarization activates sNHE. In this graph, the intracellular cAMP concentration, which influences the half activation voltage, is fixed at 1mM. The y-axis label denominator *E* is the maximal activity of sNHE. (C) SLO3 activation against pH_i_. Intracellular alkalization activates SLO3. In this graph the transmembrane potential *V*_m_ is fixed at +80mV and the phosphorylation level P_SLO3_ is kept at 0 (unincubated sperms). The y-axis label denominator 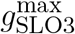 denotes 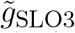 in the text. (D) An increase in pH_i_ activates SLO3 which lets potassium ions flow out and so hyperpolarizes the cell. This hyperpolarization activates sNHE which in turn extrudes protons and so increases pH_i_.

The first term on the right side of equation 1 represents the contribution of the sperm-specific sodium-proton exchanger sNHE. This contribution is always positive as the extrusion of protons has the effect of increasing the pH_i_. This transporter has a putative voltage sensor (Wang et al., 2003) and is activated by hyperpolarization in sea urchin sperm (Windler et al., 2018) (figure 1B). Moreover, the half maximal activation’s voltage of sNHE, 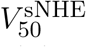, depends on the intracellular cAMP concentration. As the cAMP_i_ production by the soluble adenylyl cyclase (sAC) is a function of the concentration of 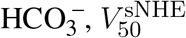 can be modeled as a Hill function as follow:

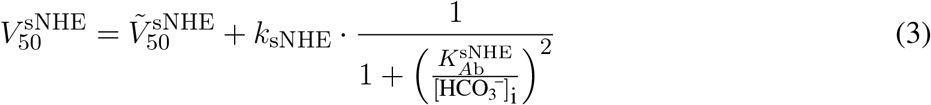

The constant 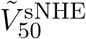 is the half maximal activation’s voltage of sNHE in absence of bicarbonate. The maximal activation shift of sNHE, *k*_sNHE_, has been set to the value of 14mV, corresponding to experimental data where [cAMP]_i_=1mM, which is well above the physiological level (Jansen et al., 2015; Garbers et al., 1982). Precise data for the voltage activation parameters of sNHE against 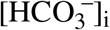 is not available in the literature; yet, based on the percentage of hyperpolarized sperm against the bicarbonate concentrations obtained by Escoffier et al. (2015) showing a jump around or below 7mM, it is reasonable to take 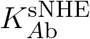 at 3mM. Finally, the pH_i_’s dependence of the activity of sNHE is taken into account as follow:

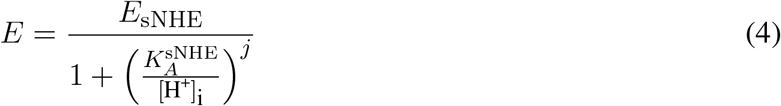

The maximal activity of sNHE (*E*_sNHE_), 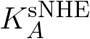and *j*, for which data is not yet available for mouse sperm, have been set to the values obtained for sodium-protons’ exchangers in fibroblasts (Boron, 2004). All parameters are given in table 1 in the appendix.

**Table 1.** Parameters values used in the simulations of the model. The references used in this table are the following: [1]:Windler et al. (2018), [2]:Jansen et al. (2015), [3]:Garbers et al. (1982), [4]:Yang et al. (2011), [5]:Zeng et al. (2015), [6]:Kirichok et al. (2006), [7]:Yeung et al. (2002), [8]:Chávez et al. (2013), [9]:Escoffier et al. (2015), [10]:Magid and Turbeck (1968), [11]:Boron (2004), [12]:Putnam (2012) *Data obtained for other cell types than mouse spermatozoa, where the values have been adjusted for the sperm’s cytosolic volume.

| Parameters |  | Value | References |
| --- | --- | --- | --- |
| $J_H^{\text{Metabolism}}$ | acid loading from metabolism | 5 $\mu\text{M/L}$ | free parameter |
| $E_{\text{sNHE}}$ | sNHE activity (for the whole membrane) | 4 mM/s | [11]* |
| $K_A^{\text{sNHE}}, j$ | Hill parameters for sNHE's $\text{pH}_i$ activation | 0.3 $\mu\text{M}$, 4 | [11]* |
| $\tilde{V}_{50}^{\text{sNHE}}, s$ | sNHE voltage activation parameters | -70 mV , 4 mV | [1] |
| $k_{\text{sNHE}}$ | sNHE's maximal voltage activation's shift | 14 mV | [1,2,3] |
| $K_{\text{Ab}}^{\text{sNHE}}$ | Hill parameter for sNHE's $\text{HCO}_3^-$ activation | 3 mM | free parameter |
| $K_{\text{Ab}}^{\text{SLO3}}$ | Hill parameter for SLO3's $\text{HCO}_3^-$ activation | 5 mM | free parameter |
| $K_A^{\text{SLO3}}, q$ | Hill parameters for SLO3's $\text{pH}_i$ activation | 32 nM , 1.65 | [4] |
| $\alpha$ | relative increase of $g_{\text{SLO3}}$ when phosphorylated | 8.3 | [8,9] |
| $\tilde{g}_{\text{SLO3}}^{\text{max}}$ | max unphosphorylated SLO3's conductance | 6.4 nS | [5] |
| $V_{50}^{\text{SLO3}}, s_{\text{SLO3}}$ | SLO3's Boltzmann fit parameters | 12.8 mV , 76 mV | [5] |
| $g_l, V_{\text{leak}}$ | leak current parameters | 1 nS , -40 mV | [8,9] |
| $k_{\text{bicar}}, K_A^{\text{bicar}}$ | Hill parameters for $\text{HCO}_3^-$ transport | $\frac{1}{3} \text{ s}^{-1}$ , 1 $\mu\text{M}$ | [11]* |
| $B_I$ | total intrinsic protons buffer | $10^4$ | [11]* |
| $P_H$ | membrane's permeability to protons | $0.005 \text{ cm s}^{-1}$ | [12]* |
| $C_m, S$ | sperm membrane's capacity and surface | 2.5 pF , 250 $\mu\text{m}^2$ | [6] |
| $V$ | sperm's cytosolic volume | 30 fL | [7] |
| $K_h$ | $\text{CO}_2$'s hydration equilibrium constant | 0.003 | [10] |

The other contributions to the variation in pH_i_ of the sperm cell are the following:

Firstly the contribution from the flux of bicarbonates, 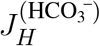, representing the sum of the fluxes through all the bicarbonate transporters. To date, the bicarbonate transporters reported in sperm are SLC26A3 (Wang et al., 2021), SLC26A6 and a putative electroneutral anion exchanger (Chávez et al., 2012), possibly a 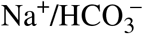 cotransporter (Vyklicka and Lishko, 2020), and the cystic fibrosis transmembrane regulator (CFTR) (Xu et al., 2007; Hernández-González et al., 2007; Escoffier et al., 2012). This bicarbonate flux, including both active and passive transports, is reduced to its most simple expression of a linear dependence on the transmembrane gradient of the bicarbonate concentration 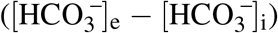, as shown in equation (7) in appendix. We made this choice as we are building a minimal model of sperm capacitation; only the core process of the feedback loop, comprising SLO3 and sNHE, is treated to produce the significant shifts in *V*_m_ and pH_i_ of the cell, regardless of details in the bicarbonate transports. Secondly, a term of passive flux of protons across the membrane 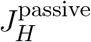, described by the Goldman-Hodgkin-Katz equation for electrodiffusion (Equation (8) in appendix), for which the permeability is here chosen in the physiological range (Putnam, 2012). This term was added to the model in order to prevent a shift to extra-low pH_i_ values when 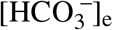 is abruptly changed.

The next contribution to the variations in pH_i_ comes from the effect of changes in extracellular carbon dioxide concentration which are instantly followed by the same intracellular carbon dioxide concentration changes as the permeability of the plasma membrane to CO_2_ is high. So in the model, the CO_2_ concentration is the same on both sides of the sperm membrane and an increase CO_2_ results in acidification of the cell as the CO_2_ is converted to bicarbonates and protons. This intracellular equilibration between carbon dioxide and bicarbonate is fast as the carbonic anhydrases (CAII, see Wandernoth et al. (2015)) catalyse the reversible reaction, as schematized in figure 1A. In the extracellular medium, as the incubation medium is devoid of carbonic anhydrases as it is usually the case *in vitro* incubations, the equilibration should be considered not instantaneous, but for simplicity we do not include this delay in our model. We nevertheless carried out simulations using explicitly the reaction rates of the conversion of CO_2_ into protons and checked that the results and conclusions presented in this paper are not different (data not shown).

The last contribution to the variations in pH_i_ is the acid loading resulting from the metabolism, taken as a constant. This acid loading, which allows the sperm to reach an equilibrium state by compensating the pH_i_ increase due to the the protons’ extruders, includes the contribution of the leak of protons from the acrosome which is an acidic organelle that has been shown to alkalize during capacitation (Nakanishi et al., 2001). The value of this parameter is adjusted so that the pH_i_ of the sperm cell in uncapacitating conditions lies in the range of the experimental data of the literature, between 6.4 and 6.85 (Zeng et al., 1996; Carlson et al., 2007; Chávez et al., 2019).

The first term on the right side of the equation (2) is the contribution of SLO3, considered with its auxiliary subunit LRRC52 (Yang et al., 2011), to the transmembrane electrical potential. It is activated by alkalization as shown by the Hill’s [H^+^]_i_ dependence of its conductance. The H^+^_i_ concentration for half occupation of SLO3 (written 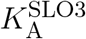) and the cooperativity coefficient q of this Hill’s function have been determined by Yang et al. (2011) using heterologous expression of SLO3 and LRRC52 in oocytes, and the corresponding activation’s curve is drawn on figure 1C for uncapacitated sperms.

The factor 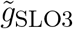 appearing in the contribution from SLO3 includes a factor representing the phosphorylation by the cSrc kinase which occurs downstream of the activation of the soluble adenylyl cyclase by intracellular bicarbonate, as follows:

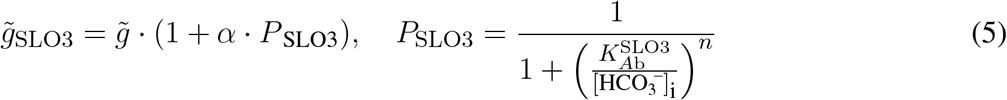

where P_SLO3_ is the phosphorylation level of SLO3 and is chosen as a Hill equation. The pathway to SLO3 phosphorylation is initiated by the activation of the soluble adenylyl cyclase (sAC) from an increase in 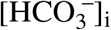, which is followed by the activation of the protein kinase A (PKA) and finally leads to the auto-phosphorylation of the cSrc kinase (Stival et al., 2015) which then phosphorylates tyrosine residues of SLO3. In fact, the phosphorylation of SLO3 is slow, of a timescale of 15min (Stival et al., 2015), but we do not include this delay in our equations because we will focus on the results concerning the steady states of the sperm cell that are reached during incubations in various media. The half maximal concentration 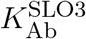 is set at a value around 5mM as shown in table 1 in appendix, and the value of the cooperativity coefficient n is set at 6, as explained in appendix. Finally the SLO3’s conductance depends not only on pH_i_ but also on *V*_m_ (Zeng et al., 2015) and this is taken into account in the factor 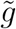 appearing in the SLO3’s conductance, independently of the pH_i_ activation, as follows:

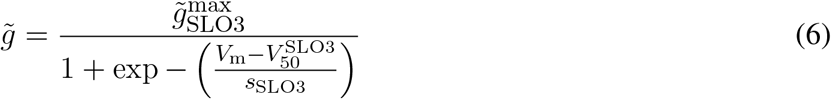

where the Boltzmann activation parameters were measured by Zeng et al. (2015) with electrophysiological techniques on spermatozoa at a pH_i_ of 8. This way of modeling the voltage activation of SLO3 independently of the intracellular pH value is based on experimental activation curves obtained by Yang et al. (2011) for various membrane voltages that indicate a pH_i_-independent voltage activation’s behavior.

The second term on the right of equation (2) is regrouping into a leak current the contribution from all other electrogenic transporters of the sperm cell membrane, where the resultant leak conductance *g*_l_ is a constant (the value of which is described in appendix). This choice of taking solely SLO3 as the effector of *V*_m_ changes during capacitation is due to the fact that we are considering the core model of a feedback loop in order to build a minimal model of capacitation. This choice is supported by the demonstration that the variation in SLO3’s conductance is responsible for the hyperpolarization of the sperm cell during capacitation (Santi et al., 2010; Chávez et al., 2013). The value of *V*_leak_ is set at 40 mV according to the transmembrane voltage obtained by Chávez et al. (2013) for SLO3’s knock-out and SLO3’s inhibited mouse sperms, both giving a value around −40 mV.

With this minimal model built around the two actors SLO3 and sNHE, we made simulations of the time evolution of the state of a sperm cell in order to check if it gives account of the capacitation process and if a positive feedback between sNHE and SLO3 could be at the core of the process. We also simulated the effects of the inhibition of SLO3 or sNHE on the state of the sperm.

The simulations, using the evolution equations (1) and (2), were performed using the software XPPAUT 6.11 (Free Software Foundation Inc., Cambridge, USA). In all simulations, the extracellular pH (pH_e_) is fixed at the value of 7.4 and the temperature is set at 37 °C, as usually the case in sperm incubating media. Source code is available upon request.

## 3 RESULTS

### 3.1 The positive feedback between SLO3 and sNHE causes capacitation

In order to find if the model reproduces the transition from a depolarized acidic state (uncapacitated) to a hyperpolarized alkaline state (capacitated) of the sperm, we performed a simulation of the incubation of a sperm cell as it is usually done in the laboratory, consisting of the incubation into a solution containing 15mM bicarbonate 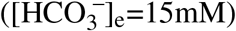.

This protocol is represented on the top graph of figure 2A showing a step of 15mM of bicarbonate at time t=2min. The initial concentration of bicarbonate in the medium is set at 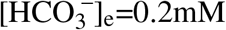, which corresponds to a solution of pH=7.4 at equilibrium with ambient atmosphere. As indicated in the graphs of the pH_i_(t) and *V*_m_(t) on figure 2A, before the bicarbonate step (t<2min) the sperm is at a steady state of (pH_i_,*V*_m_)≃(6.6,−45 mV). After the 15mM pulse of bicarbonate at time t=2min, the state of the sperm shifts to a pH_i_ of 7.1 and a *V*_m_ of -75mV. This transition of the sperm’s state is here fast, of the timescale of 5 minutes; however, adding the phosphorylation delay of SLO3 results in a transition’s timescale of 15 minutes (data not shown), in accordance with data of the literature for capacitation of mouse sperm cells (Stival et al., 2015; Arnoult et al., 1999).

**Figure 2.**
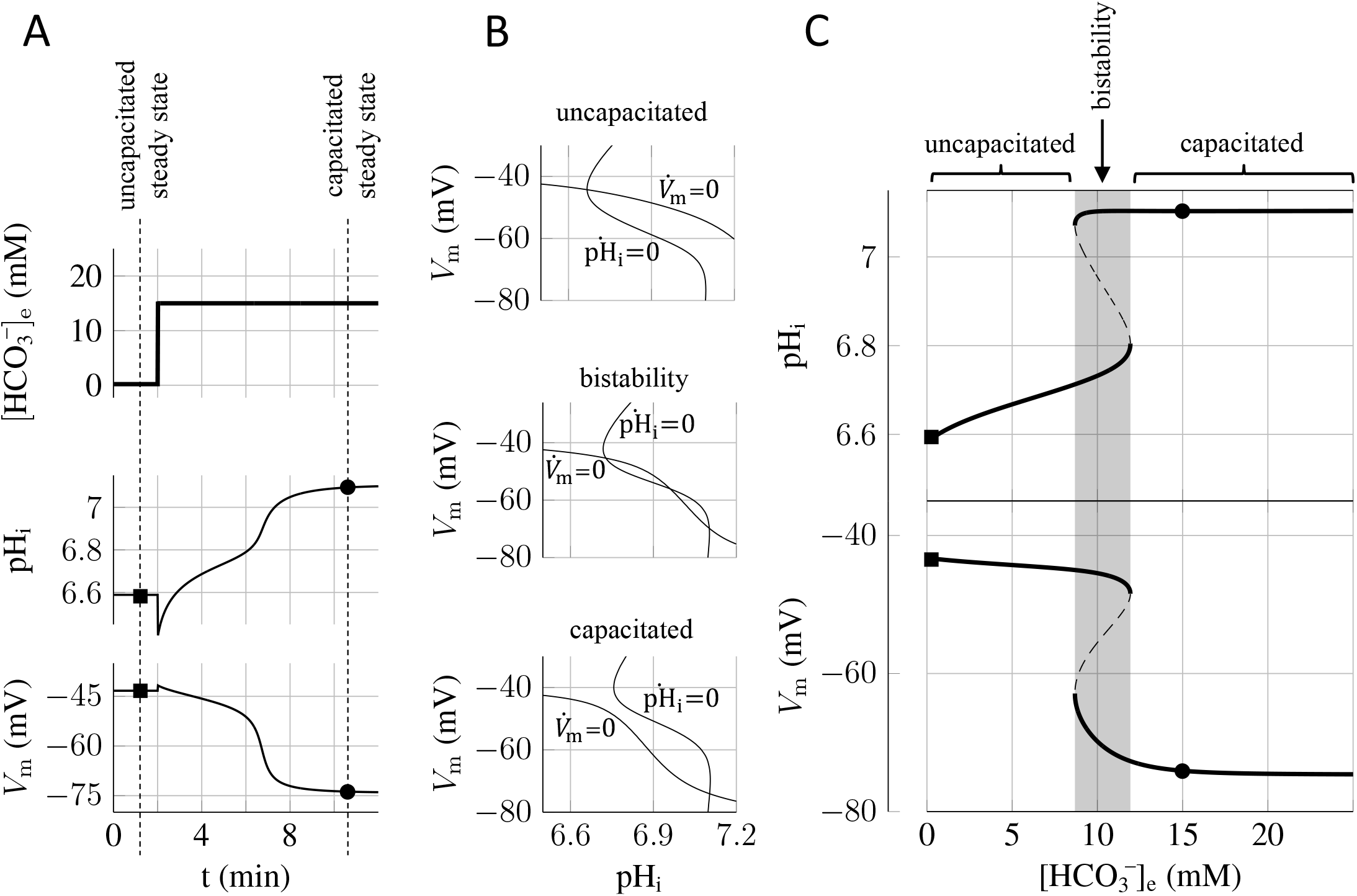
Transition between uncapacitated and capacitated state as the bicarbonate concentration is increased. (A) Time evolution of the pH_i_ and *V*_m_ following a step from 0.2mM to 15mM extracellular bicarbonate 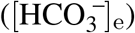 at time t=2min. Following the increase in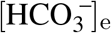, the pH_i_ rises from the initial uncapacitated value of around 6.6 to the capacitating value above 7.1. This significant increase comes with the cell’s hyperpolarization from − 45 mV to − 75 mV and reflects the feedback loop between SLO3 and sNHE. Beware the instant phosphorylation of SLO3 used in the simulations induces a transition in the five minutes’ time scale, faster than the 15 minutes timescale necessary for the hyperpolarization of mouse sperm. (B) Graphs of the nullclines of the mathematical model for three different extracellular bicarbonate concentrations. Equalling to zero the two equations (1) and (2) leaves us with two nonlinear equations of the two variables pH_i_ and *V*_m_, providing two lines (called nullclines) in the (pH_i_,*V*_m_) plane as shown on each three graphs. The lines labeled 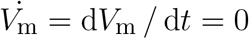 correspond to the states for which the transmembrane potential does not change with time, and the lines labeled 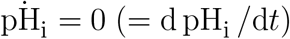 correspond to the states for which the cytosolic pH of the sperm does not change with time. The intersections of these two lines give the steady states of the sperm cell at which both *V*_m_ and pH_i_ do not change in time. The graph on top shows the two nullclines for 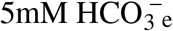; middle graph: 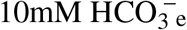; bottom graph: 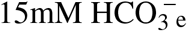. (C) Bifurcation diagrams showing the steady states of the sperm as a function of the concentration of 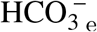. The upper diagram shows the pH_i_ of the steady states and the lower diagram shows the transmembrane potential *V*_m_ of the steady states. On the left of the gray region appear in thick solid lines the stable steady states corresponding to an uncapacitated sperm. On the right of the gray region (i.e. for high 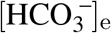 appear the stable steady states (thick solid line) corresponding the capacitated sperm. The transition between the uncapacitated state and the capacitated state occurs in the gray region where two stable steady states and one unstable steady state (represented in dashed line) are found. This gray region corresponds to the values of 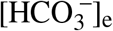 where the system is bistable: the sperm can be found in two different stable steady states.

This simulation of the time evolution of the state of the sperm therefore shows two steady states, one at 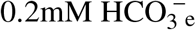and one at 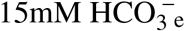, which are reported on the two diagrams of figure 2C (black squares and bullets), one corresponding to the uncapacitated state of the sperm and one corresponding to the capacitated state of the sperm.

### 3.2 Bistability of the capacitation process

Having found two steady states of the sperm, one at 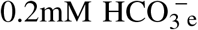 and one at 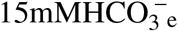, we wondered if these sperm states were the only possible ones for each of these extracellular bicarbonate concentration, and whether there is a threshold in 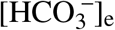 above which the uncapacitated sperm will spontaneously shift to the capacitated state of low *V*_m_ and high pH_i_. As a positive feedback loop can give rise to bistability we can expect that for some values of the parameter 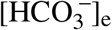 we would find two possible stable states, one capacitated and one uncapacitated.

The top graph of figure 2B shows that when 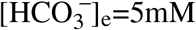 there is only one possible steady state for the sperm. When 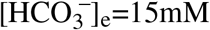 (bottom graph of figure 2B), there is also only one steady state, at the point (pH_i_,*V*_m_)≃(7.1, − 75 mV). This latter state is hyperpolarized and alkaline, and so corresponds to a sperm’s capacitated state, as the one found in figure 2A where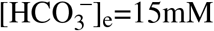.

Between these two bicarbonate concentrations of 5mM and 15mM however, at 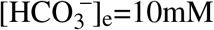 we find three intersections of the nullclines, which are shown in the middle graph of figure 2B, and indicating that the sperm can be found in three different steady states, one of these three states corresponding to a capacitated state of the sperm at *V*_m_ ≃ − 70 mV and pH_i_ ≃7.1. In order to gain insight of what is happening here we computed the steady states for all values of 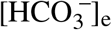and plotted these steady states in two diagrams shown on figure 2C, where we observe that a transition between uncapacitated states and capacitated states happens when the extracellular bicarbonate concentration crosses a threshold at around 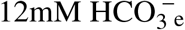. Below 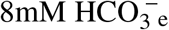are found the uncapacitated states of the sperm and above 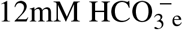 are found the capacitated states of the sperm, and between these two appears a region (shaded in the figure) where three steady states of the sperm coexist. The analysis of the dynamics of the system shows that the steady states in the middle (dashed lines in figure 2C) are unstable; in reality the sperm will never be found at such steady states because any imperceptible fluctuation in the physiological conditions brings the sperm state away from these unstable steady states. Therefore, in the shaded region, the sperm can be found in two different steady states; it’s a phenomenon of bistability where the state of the sperms will depend on what happened before they were brought to this bistable region. Indeed, if we start on the left of the diagram where 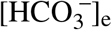is low, and increase slowly the medium’s bicarbonate concentration to the bistable region, the sperm will be found at the state (pH_i_,*V*_m_) ≃(6.7, −45 mV), but if the sperm is prepared on the right of the diagram, decreasing the 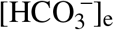 to the gray region will keep the sperm at the capacitated state near (7.1, − 70 mV).

### 3.3 The capacitation switch has hysteresis yet is reversible

The bistability arising from the feedback loop between sNHE and SLO3 implies that the sperm will chose one of the two stable states according to its initial state. This hysteresis phenomenon implies that once the sperm has chosen one of the two stable states, it will remain locked in this state regardless of small changes in the surrounding physiological conditions.

That is what we show here with a simulation of an experiment where a hysteresis effect resulting from this bistability arises in the state of the sperm. The simulation, shown on figure 3, consists in preparing the sperm cell in the bistability region at 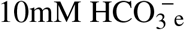before imposing a pulse of 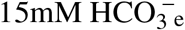 that brings the sperm to a capacitated state. We find that the capacitated state is robust as the sperm remains capacitated when the bicarbonate concentration is brought back to its initial value of 10mM after the pulse. This is a hysteresis effect and it shows that once the sperm cell is capacitated, it is blocked in this state regardless of small variations in the physiological conditions surrounding the sperm cell. We find however that this switch is reversible as a sufficient depletion of 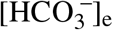 for some time is able to bring the sperm back to the uncapacitated state.

**Figure 3.**
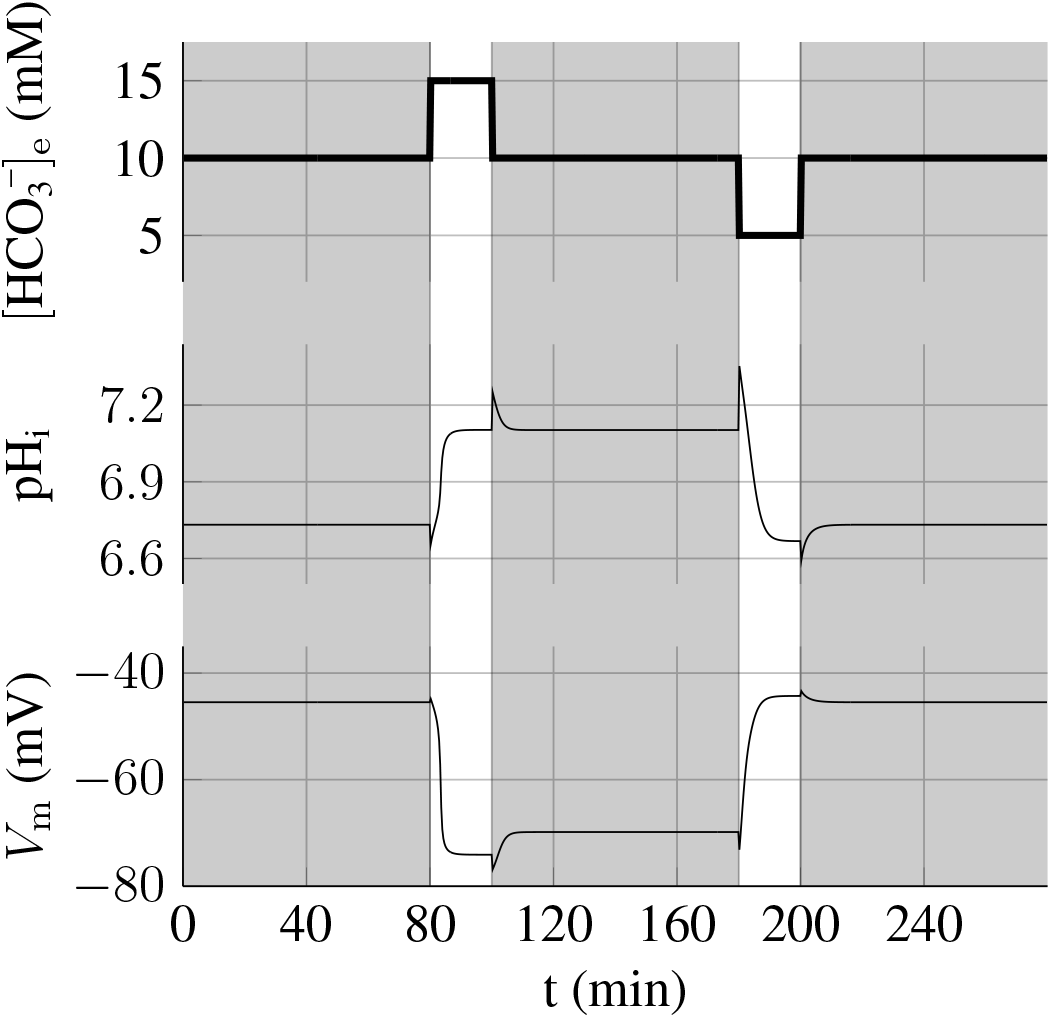
Hysteresis and reversibility of the capacitation process. After having prepared the sperm at 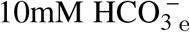, a pulse of 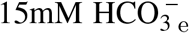 at t=80min switches the sperm to the capacitated state (i.e. *V* ≃-70mV and pH_i_ 7.1). However, when 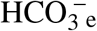 is brought back to the initial value of 10mM, the sperm remains in the capacitated state. Nevertheless, when 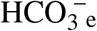 is pulsed down to 5mM for 20min, the state switches back to the initial non-capacitated state. The extracellular pH is always fixed at 7.4.

### 3.4 SLO3’s inhibition is more robust than sNHE at preventing capacitation

The proposed minimal model can also be used as a tool in order to predict how the capacitation process can be blocked efficiently. We simulated the time evolution of the sperm’s states when either SLO3 or sNHE is inhibited and have examined the transition between the uncapacitated and capacitated states. The result of the simulation of the incubation of sperm in 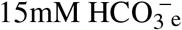 when SLO3 is inhibited at a level of 70% is presented on figure 4A and shows that capacitation does not occur anymore; the sperm does not hyperpolarize and the intracellular pH shows an increase of only 0.2 unit.

**Figure 4.**
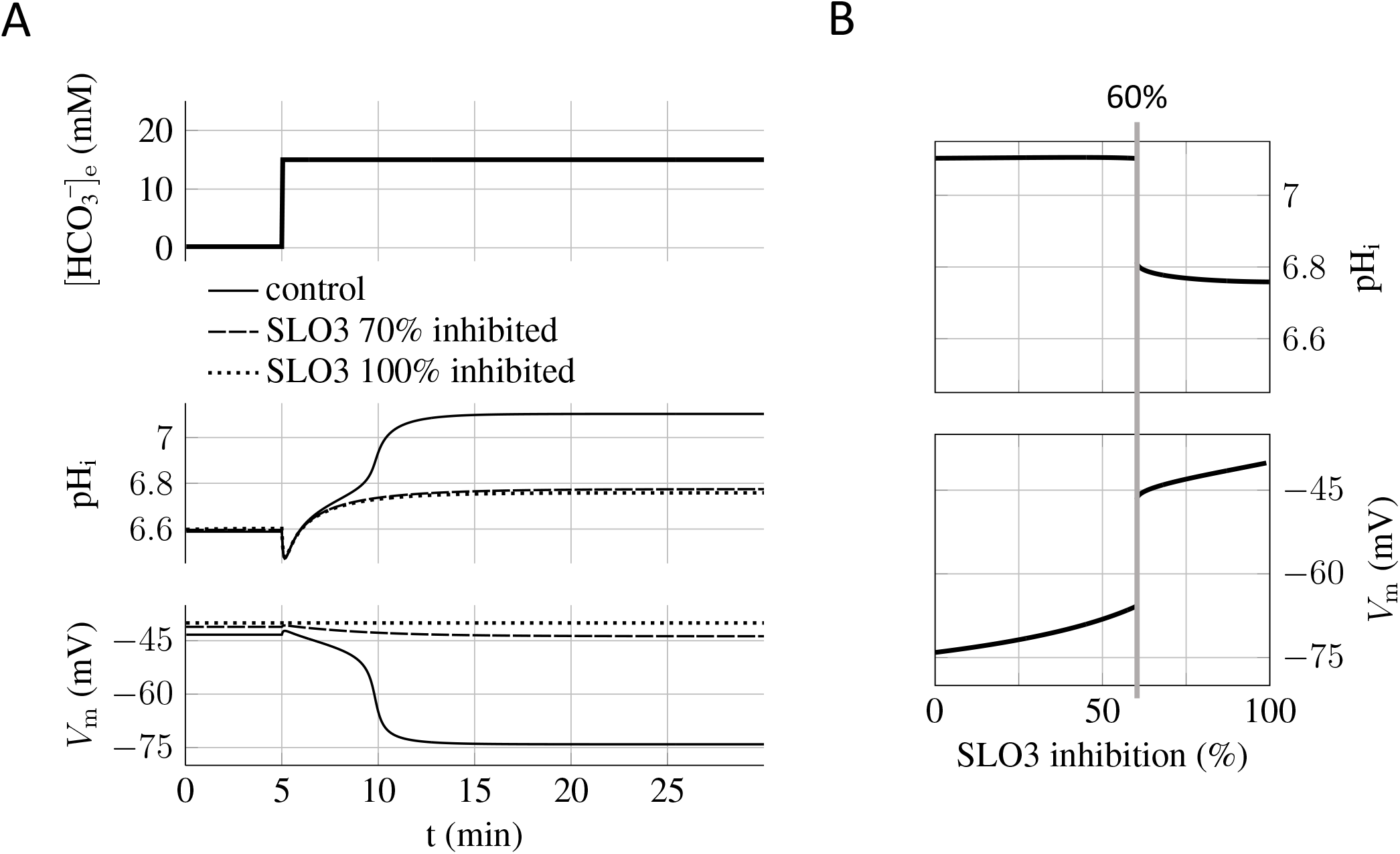
Capacitation dependence on SLO3 inhibition (A) When SLO3 is 70% inhibited, the transition to the capacitated state does not occur and the state is blocked at the values of pH_i_=6.8 and *V*_m_=-45mV. In the case of 100% inhibition of SLO3, the model shows no variation in *V*_m_ during capacitation. (B) Graph of the steady states of the sperm against the percentage of SLO3 inhibition. These are the steady states reached after the incubation in capacitating medium (15mM bicarbonate) of sperms previously prepared at 0.2mM bicarbonate. A threshold appears at 60% inhibition above which both the increases in pH_i_ and hyperpolarization are much reduced.

By carrying on the same simulations with different inhibition percentages of SLO3 we find a threshold at 60% above which capacitation is prevented (figure 4B); the pH_i_ will increase no more than 0.2 unit, from 6.6 to less than 6.8, and the transmembrane potential will not hyperpolarize below −45 mV. The existence of this inhibition threshold above which the sperm does not capacitate anymore is due to the fact that the system is brought into the bistable region and will therefore remain blocked to the uncapacitated state during incubation.

When simulating an incubation at the same concentration of 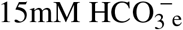 but with 70% inhibition of sNHE, the effect on capacitation is similar to the SLO3 inhibition (data not shown), but when increasing the 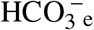 concentration of the incubation medium we find that the sNHE inhibition is not as effective at preventing the capacitation switch as the inhibition of SLO3. This is illustrated on figure 5 showing the states of incubated sperms as a function of the 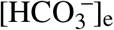 at a fixed 95% inhibition. The SLO3 inhibition keeps the state depolarized for all bicarbonate concentrations (figure 5A) but in contrast, when 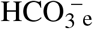 crosses the threshold of 20mM, the capacitation switch is not prevented by a 95% sNHE inhibition (figure 5B). The prevention of the hyperpolarization in the case of the SLO3’s inhibition is due to the fact that its conductance remains small compared with the leak conductance.

**Figure 5.**
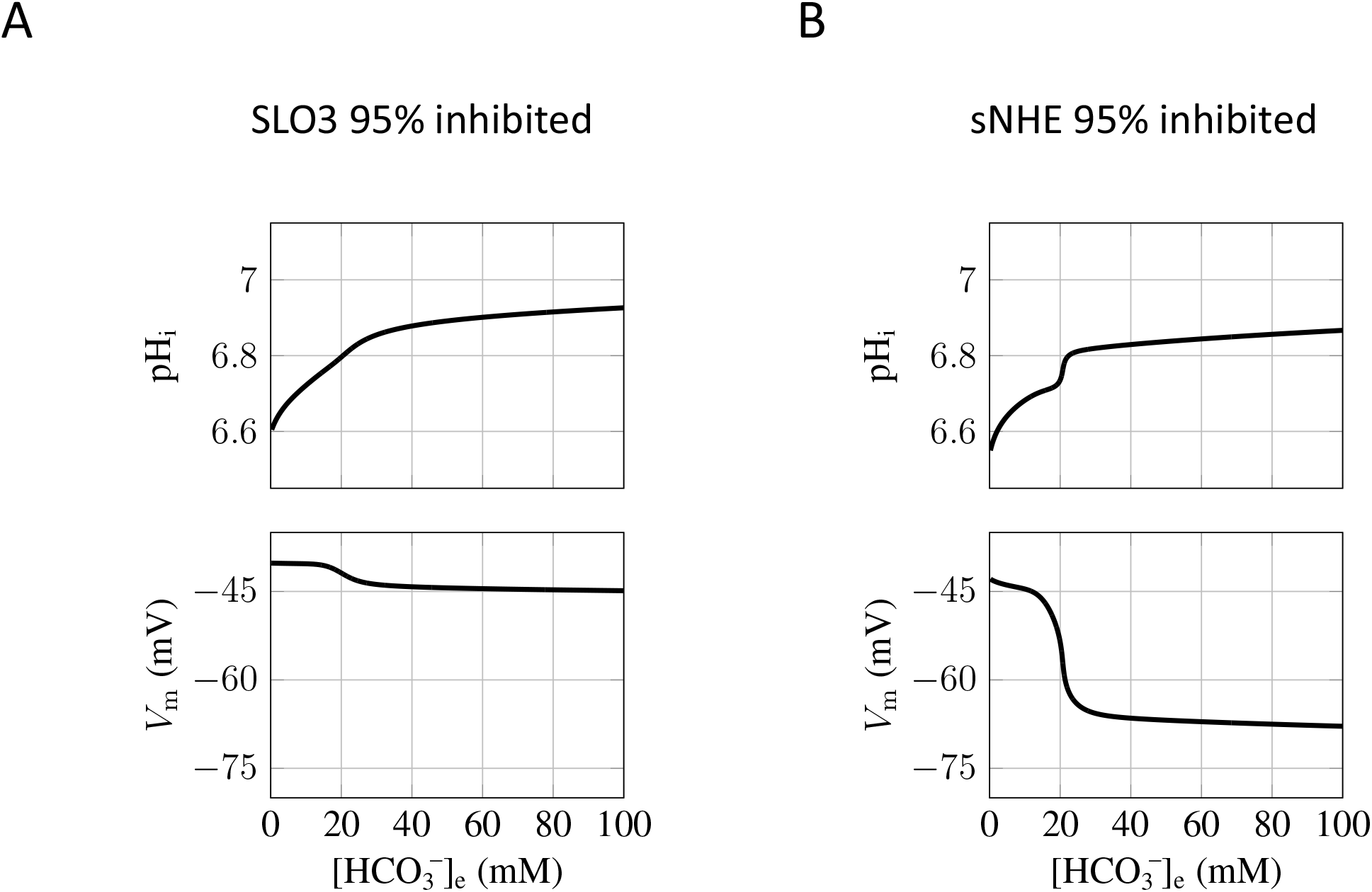
SLO3 inhibition is more robust than sNHE at preventing capacitation. (A) When SLO3 is 95% inhibited, the switch to the capacitated sperm does not occur for any 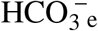 concentration. (B) When sNHE is 95% inhibited, the capacitation switch is not prevented when the 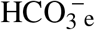 concentration exceeds 20mM.

## 4 DISCUSSION

In this work we have built a minimal model for the capacitation of mouse spermatozoa as a function of the incubating medium’s bicarbonate concentration. We modeled the sperm’s state evolution on the minimal basis of the effect of sNHE and SLO3 on pH_i_ and *V*_m_. By doing so, for the homeostasis of sperm pH_i_ we included, in addition to sNHE and SLO3, the alkalizing effect of bicarbonate transporters and the acidifying effect of the metabolism of the sperm cell.

The majority of the parameters values used in the model are based on existing data, with the exception of three which remain free. These free parameters are the acid loading from the metabolism and the bicarbonate concentration for downstream half activation of sNHE or SLO3. Nevertheless, the acid loading from the metabolism, which compensates the alkalizing effects of the protons and bicarbonate transporters in order to allow pH homeostasis, is inherently physiological as the alkalizing parameters are physiological.

Concerning the half activation of sNHE and SLO3 downstream of the protein kinase A activation by bicarbonate, our parameter’s values are taken consistently with results from Stival et al. (2015) giving the percentage of hyperpolarized sperms as a function of the 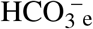 concentration, which indicates a threshold below 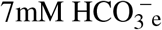 above which a substantial proportion of sperms do hyperpolarize during the incubation. However, the exact value of these two parameters do not affect the essence of the feedback loop and bistability in our model; it rather shifts the shaded bistability region (shown in figure 2C) to the right or to the left. In particular, decreasing 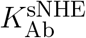 will shift the capacitation threshold from around 12mM (as in figure 2C) to a lower concentration (data not shown).

Our minimal model gives account of the experimentally observed hyperpolarization and alkalization of the mouse sperm’s cell during capacitation. Indeed the hyperpolarization of the sperm’s cell to -75mV lies in the range (*V*_m_<70mV) found in the literature for capacitated sperms (Arnoult et al., 1999; Escoffier et al., 2015). We emphasize here that our results concern individual spermatozoa that effectively capacitate, and not a “capacitated population” of sperms that consists of capacitated and uncapacitated subpopulations for which the measured hyperpolarization represents the sum of the depolarized uncapacitated subpopulation and the hyperpolarized capacitated subpopulation. In such heterogenous populations, the measured hyperpolarization is around -60mV (Arnoult et al., 1999; Stival et al., 2015).

Concerning the evolution of the sperm’s pH_i_ during capacitation, our model shows an increase in pH_i_ from 6.6 before incubation to 7.1 after incubation. The uncapacitated value given by the model was in fact adjusted to the value of 6.6 obtained by Chávez et al. (2019), and the capacitated value of 7.1 is in the range around 7.2 obtained recently by Ferreira et al. (2021). However, we would like to stress here that our model’s pH_i_ values of 6.6 before capacitation and 7.1 after capacitation are not fixed and do depend on the exact value of the acid load from the metabolism which is a free parameters as mentioned here above.

Our analysis of the sperm’s steady states as a function of the bicarbonate concentration in the incubating medium showed the existence of a bistability region where the sperm can be found in two different states and we showed that once the sperm has reached the hyperpolarized and alkaline state of capacitated sperm, it will remain capacitated regardless of small fluctuations in the bicarbonate concentration. In addition, we show that this hysteresis phenomenon in the cellular response allows the encoding of transient signals, like a pulse in the external bicarbonate concentration, by long-lasting changes both in pH_i_ and *V*_m_. This capacitation switch can be reversed back as a sufficiently strong decrease in 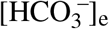 will bring the sperm back to an uncapacitating state. The existence of this hysteresis phenomenon resulting in a capacitation threshold which is different from the decapacitation threshold is a prediction of the model that can be tested experimentally. On a functional level, this predicted shift in bicarbonate sensitivity allows the capacitation process to be more robust in presence of local variations of bicarbonate concentration, ensuring that, once capacitated, sperm cells are less sensitive to these fluctuations.

Concerning the inhibition of capacitation, when the sperm is incubated in a 15mM bicarbonate solution we find a switch at 60% inhibition of SLO3 above which capacitation is prevented. Even though the inhibition of sNHE has similar effect on capacitation in an incubation medium of 15mM bicarbonate, we find that inhibiting SLO3 is more effective than inhibiting sNHE when the bicarbonate concentration is increased. Indeed, a 95% inhibition of SLO3 prevents capacitation for all possible bicarbonate concentrations that could be found in the genital tract of the mammals (Maas et al., 1977), which is not the case when instead sNHE is 95% inhibited. This stronger effect of SLO3 inhibition on capacitation is not surprising as we may expect inhibition of ion channels to have a much larger impact on membrane potential than the inhibition of a carrier system like sNHE, which has a much lower turnover rate for ion transport. It appears therefore that the inhibition of SLO3 is a better candidate for the development of non-hormonal male contraception.

The human sperm however is different from the mouse as hSLO3 is strongly activated by calcium and less by pH (Brenker et al., 2014). Our model could be adapted and applied to human sperm by adjusting the activation curves of SLO3 and taking into account the voltage-gated hydrogen channel 1 which has been shown to be the dominant proton conductance in human sperm (Lishko et al., 2010).

The proposed minimal model has allowed us to identify the core molecular mechanism which is likely to control murine sperm capacitation. It may be extended to include calcium dynamics and further refined to the human case. Concerning the increase in intracellular calcium during capacitation, the feedback loop between sNHE and SLO3, which robustly switches the state of the sperm to an elevated intracellular pH, is likely to keep the pH_i_-sensitive Catsper channel activated during capacitation, and moreover, the reversibility of the switch implies that this capacitation’s aspect of calcium influx can be reversed back. In addition, as we have modeled the soluble adenylyl cyclase activity only through the increase in intracellular bicarbonate concentration and not by increases in intracellular calcium concentration which are known to activate this adenylyl cyclase (Litvin et al., 2003). An extended version of the model, explicitly including Catsper, could take into account this additional feedback. Moreover, in the case of human sperm capacitation, the activation of hSLO3 by calcium should also be included in future developments of our model.

To conclude, using a modeling approach, we have identified a possible core molecular mechanism underlying the transition between the uncapacitated and capacitated state in mouse sperm and studied its dynamic as well as identified pharmacological targets regulating this process. These results may be relevant in the possible development of treatments of male infertility and non-hormonal male contraception, as the inhibition or stimulation of these actors of sperm’s capacitation can modulate the ability of sperm to fertilize the egg. For this, the proposed model should be extended to the human case.

## APPENDIX

In this section are presented the complete set of functions and equations of the model in addition to the main equations (1) and (2). A table of the values of the parameters used in the simulations of the model is also given.

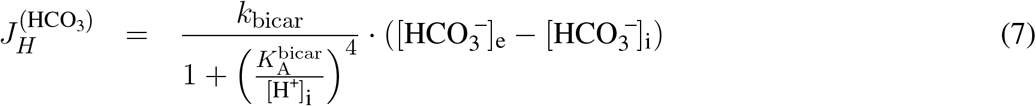

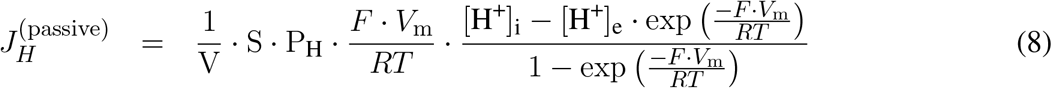

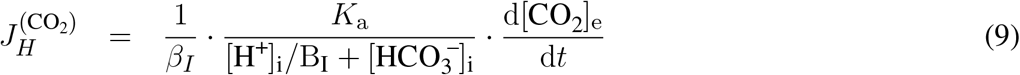

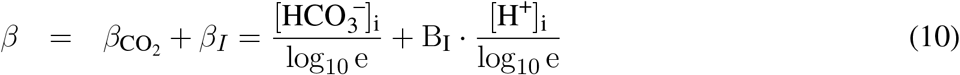

Concerning the bicarbonate dynamics, the following equilibrium equation is used, both for extracellular and intracellular medium concentrations:

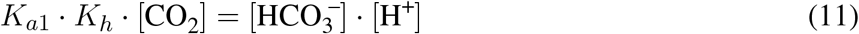

The value of the α factor appearing in equation 5 and representing the relative increase of the SLO3’s conductance when phosphorylated, is fixed by assuming that the pH_i_ and *V*_m_ of the capacitated sperms is 7.1 and -80mV respectively. Indeed, when using these latter values with the results of potassium permeabilities before and after capacitation of sperm populations obtained by Chávez et al. (2013), together with the percentage of sperms that effectively do capacitate in the populations subjected to capacitating conditions (Escoffier et al., 2015) and with the values of the pH_i_ before capacitation found in Chávez et al. (2019), our description of SLO3’s conductance predicts then α = 8.3.

We then set the value of g_*l*_ to 1 so that the model accounts for the sperm’s transmembrane voltage shift from around -40mV to around -80mV during capacitation (Chávez et al., 2013).

Finally the value of the cooperativity coefficient *n* for the SLO3 activation by 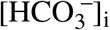 (equation 5) has been set at the value of 6 in order for the evolution *V*_m_(t) predicted by the model (including here the phosphorylation delay) to reproduce the results of the time evolution of *V*_m_ obtained by Stival et al. (2015) which show a hyperpolarization occurring between 15 minutes and 30 minutes of incubation in a 15mM 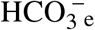 capacitating medium (data not shown).

Other common constants used in the model at the temperature of the simulations T=310K in physiological solutions: *V*_K_= − 84.9 mV, F=9.65 × 10^4^ C mol^− 1^, *R*=8.31 J mol^−1^ K−^1^, K_*a*1_=2.8 × 10^4^ M (carbonic acid’s first dissociation constant).

## AUTHOR CONTRIBUTIONS

B. de Prelle, P. Lybaert and D. Gall designed the research, carried out the simulations and wrote the article.

## ACKNOWLEDGMENTS

We thank Genevieve Dupont and Celia M. Santi for their remarks and suggestions during the construction of the model.

